# Multiple non-auditory cortical regions innervate the auditory midbrain

**DOI:** 10.1101/627232

**Authors:** Bas MJ Olthof, Adrian Rees, Sarah E Gartside

## Abstract

Our perceptual experience of sound depends on the integration of multiple sensory and cognitive domains, but the networks sub-serving this integration are unclear. There are connections linking different cortical domains, however we do not know if there are also connections between multiple cortical domains and subcortical structures. Retrograde tracing in rats revealed that the inferior colliculus – the auditory midbrain - receives dense descending projections from not only the auditory cortex, but also the visual, somatosensory, motor, and prefrontal cortices. While all these descending connections are bilateral, those from sensory areas show a more pronounced ipsilateral dominance than those from motor and prefrontal cortices. Anterograde tracing from cortical areas identified by retrograde tracing, showed cortical fibres terminating in all three subdivisions of the inferior colliculus, targeting both inhibitory and excitatory neurons. These findings demonstrate that auditory perception is served by a network that includes extensive descending connections from sensory, behavioural, and executive cortices.

## INTRODUCTION

Sensory processing has traditionally been viewed as the centripetal flow of information from a sense organ to a modality-specific region of cerebral cortex. This view is now being revaluated on two grounds. First, evidence for extensive feedback connections from higher levels to earlier stages in the processing hierarchy demonstrates the flow of information is not one way. Such top-down processing is in keeping with a growing appreciation that perception involves the interaction between incoming sensory information and predictions or expectations based on prior experience (Rao and Ballard, 1999; Bastos et al., 2012). A second challenge to this view of sensory processing, is evidence of multimodal responses in areas of cortex previously considered to be exclusively unimodal, including so called ‘primary’ regions (Kayser and Logothetis, 2007; Bizley and King, 2008, 2009; Bastos et al., 2012).

Top-down and cross-modal sensory interactions have been most extensively explored in the cerebral cortex, but the organisation of the auditory system, with its many subcortical centres, means that such interactions could potentially occur much earlier in the pathway. The inferior colliculus (IC) - the midbrain auditory centre-receives extensive top-down input from the auditory cortex (Diamond et al., 1969; Beyerl, 1978; Andersen et al., 1980; Saldana et al., 1996; Winer et al., 1998; Budinger et al., 2000; Bajo and Moore, 2005; Coomes et al., 2005; Bajo et al., 2007). Cortico-collicular connections are particularly evident in the dorsal and lateral cortex of the IC (ICD and ICL), but are also seen in the central nucleus (ICC) (Saldana et al., 1996), which is the main recipient of afferent input from lower level brainstem centres. Cortico-collicular connections mediate both short and long term plasticity that influences the response properties of neurons in the IC to sounds (Gao and Suga, 2000; Suga and Ma, 2003; Bajo et al., 2010; Bajo and King, 2013).

While research on cortico-collicular connections has focussed almost exclusively on those originating from the auditory cortex, one previous anatomical study, conducted before the advent of modern tracers, suggested that inputs to the IC might originate from more diverse cortical sources. Thus, Cooper and Young (1976) found degenerating fibres in the IC following lesions to the visual, ‘somaesthetic’ (somatosensory) and motor cortices, suggestive of cortico-collicular inputs from these non-auditory cortices. Surprisingly, in spite of this evidence, the intriguing possibility that the IC receives inputs from cortical areas other than those sub-serving audition, has not been systematically addressed.

Here we use retrograde and anterograde tracing to identify definitively which regions of the cerebral cortex project to the IC. We show that there are projections from sensory, motor, and executive cortices that innervate all subdivisions of the IC. These findings show that our understanding of perceptual processing, and specifically the contribution of the auditory midbrain, must be revised to take account of these several and diverse sources of top-down cortical input. We speculate that such feedback mechanisms may provide multisystem prediction signals that are essential for the identification and separation of sound sources.

## METHODS

### Animals

Experiments were performed in accordance with the terms and conditions of a license (PPL 60/3934) issued by the UK Home Office under the Animals (Scientific Procedures) Act 1986 and with the approval of the Local Ethical Review committee of Newcastle University. Male Lister-hooded rats (250-350 g) were obtained from Charles River and kept, for at least 7 days after arrival, in the Newcastle University Comparative Biology Centre under standard housing conditions, 12h light/dark cycle with food and water available ad libitum.

### Surgical procedures

Animals were heavily sedated with ketamine/medetomidine (≈15 / 0.2 mg/kg, i.p.) and the scalp was shaved. Local anaesthetic cream (Emla™) was applied to the nose and a paediatric nasogastric tube was inserted into one nostril for the delivery of isoflurane in O_2_. The concentration of isoflurane was adjusted (1-4 % in ≈0.1 litres/min O_2_) to provide a surgical plane of anaesthesia throughout. Animals were placed in a stereotaxic frame using atraumatic ear bars (David Kopf) with the tooth bar set at −0.3 mm (flat skull position). Body temperature was measured with a rectal probe and maintained at 37°C with a homeothermic blanket (Harvard Instruments). A midline incision was made in the scalp and the periosteum was removed to reveal the skull.

For **retrograde tracing** experiments (n=8), a craniotomy was performed over both ICs (co-ordinates from Bregma AP: −8.5 mm, ML: ± 2.0 mm) leaving a bridge of bone in the midline. Retrobead™ latex nanoparticles (Lumafluor) were injected using a Nanoject™ programmable injector (Drummond Scientific) fitted with a borosilicate glass capillary pipette (1.14 mm OD, 0.53 mm ID, Drummond Scientific) which had been pulled to a fine tip and broken back to allow filling from the tip. Pipettes were preloaded with aloe vera and a small volume of Retrobeads at the tip end. Red Retrobeads were injected into the right IC and green Retrobeads into the left IC. In each case the pipette was initially lowered in the ICC to a depth of −4.5 mm from the brain surface and 200 nl Retrobeads were injected over a period of 2 minutes. The pipette was left *in situ* for 2 min before being raised to −3.3 mm where a further 200 nl was injected, and then to −2.3 mm where a final 200 nl injection was made. To reduce the likelihood of the injection being drawn upwards through the tissue following the final injection, the pipette was left in place for 5 minutes before it was removed.

For **anterograde tracing** experiments, TRITC- or fluorescein-labelled dextran was injected into either one or two cortical sites (n=10). In four animals a craniotomy was performed over the right auditory cortex only (AP:−5.2, ML: 4.2-5.3 mm). A glass capillary pipette (1.14 mm OD, 0.53 mm ID, Drummond Scientific) pre-loaded with TRITC- or fluorescein-labelled dextran (10,000 m.wt, D1816 and D1821, Invitrogen, 10mg/ml in sterile 0.9 % saline) fitted to the Nanoject injector was lowered at an angle of 30-20° away from the midline). Injections (200 nl/site x 2 minutes) were made at three positions in the auditory cortex corresponding to AP: −5.2, ML: ±7.4, DV: −5.6 mm; ML: ± 6.8, DV: −4.2 mm; and ML: ±5.8, DV: −1.6 mm. In a further six animals, similar injections were made into the cortical regions as follows: 1) right prefrontal /left visual; 2) right motor only; 3) right visual /left prefrontal; 4) right prefrontal/left visual; 5) right somatosensory/left prefrontal; 6) right visual/left somatosensory. Injections were made at multiple sites in each cortical regions as follows: prefrontal cortex: AP: +3.2, ML: ±0.5, DV −4.3, −2.8, and −1.6 mm); motor cortex (AP: +3.2, ML: = 1.0, DV: −1.0 mm and ML: −1.7, DV: −2.2mm,); somatosensory cortex, (AP: +2.12, ML: ±5.0 DV −1.9 and−1.0 mm); and visual cortex (AP:−8.0, ML: ±2.5, DV: −0.8mm, and ML: ±4.07 1.30 mm; and ML: ±5.1, DV: −1.0 mm). TRITC-labelled dextran (red) was always injected into the right hemisphere and fluorescein-labelled dextran (green) was injected into the left hemisphere. Following injection, the pipette was left in place for 5 min to minimise the spread of the injection.

Towards the end of the surgical procedure, animals received an injection of meloxicam (15 μg, s.c.) to provide analgesia during recovery. When injections were complete, the scalp wound was stitched using Vicryl 4.0 suture (Ethicon) and local anaesthetic cream (Emla) was applied to the wound. Animals were allowed to recover from the anaesthetic in a warm environment and were then returned to their home cage and normal housing conditions. The next morning, a second dose of meloxicam (15 μg, s.c.) was administered to provide on-going analgesia.

### Tissue collection and processing

Between 48 and 60 hours following Retrobead injection, or 7 to 8 days following injection of dextran, animals were deeply anaesthetized with pentobarbital (≈400 mg/kg) and transcardially perfused with ≈100ml heparinised 0.1 M phosphate buffered saline (PBS, composition (mM):-NaCl: 66, Na_2_HPO_4_: 16, KH_2_PO_4_ 3.8) followed by ≈ 100 ml 4% paraformaldehyde in PBS (PFA). The brain was removed, post-fixed overnight in 4 % PFA, and then cryoprotected in 30 % sucrose. Cryoprotected brains were stored at −80 °C until sectioning.

For **retrograde tracing** experiments, the brain was sectioned on a cryostat from the cerebellum to the olfactory bulb. Coronal sections (30 or 40 μm) were collected onto gelatine subbed slides, air dried, and stored at −20 °C. Before imaging, slides were dipped in DAPI (1 μg/ml, 5 min), air dried and cover-slipped with Fluoroshield mounting medium (Sigma-Aldrich).

For **anterograde tracing** experiments, the cortical injection sites and the IC were sectioned. Coronal sections (30 μm) were cut on a rotary microtome and collected into antifreeze solution (30 % ethylene glycol, 30 % sucrose, 1 % polyvinyl pyrrolidone (PVP)-40 in PBS) (Watson et al., 1986) and stored at −20 °C. Sections from the injection sites and the IC were mounted on glass slides, dipped in DAPI (1 μg/ml, 5 min), rinsed and cover-slipped with Fluoroshield mounting medium (Sigma-Aldrich). One mid rostro-caudal IC section from each animal was immunolabelled for GABA (rabbit anti-GABA antibody 1:1000, Sigma-Aldrich Cat# A2052, RRID:AB_477652), and the neuronal marker NeuN (mouse anti-NeuN antibody 1:1000, Merck Millipore Cat# MAB377, RRID: RRID:AB_2298772). Sections were washed with PBS (3×10 min), incubated in 1 % NaBH_4_ for antigen retrieval (30 min), and washed in PBS (3×10 min) before being incubated (4 h at room temperature followed by overnight at 8 °C) in a mixture containing the primary antibodies made up in a block buffer comprising 1 % bovine serum albumin (Sigma-Aldrich), 0.1 % porcine gelatine (BDH), 50 mM glycine (Fisher) in PBS. Sections were washed again (3×10 min PBS) before incubation (2 h at room temperature) with a mixture of AlexaFluor 488 or 568 goat anti-rabbit secondary (1:500, Life Technologies) and biotinylated horse anti-mouse secondary (1:500, Vector labs) in 5 % normal goat serum (Sigma-Aldrich) in PBS. Sections were washed in PBS (3×10 min) and incubated with Cy5 streptavidin (1:500, Life Technologies) in PBS (1 h at room temperature). Finally sections were washed in PBS (10 min), incubated in DAPI (1 μg/ml, 10 min), washed again in PBS (10 min) and dipped in distilled water before mounting on plain glass slides. Slides were cover-slipped with Fluoroshield (Sigma-Aldrich).

### Imaging

For **retrograde tracing** experiments, first the injection sites were examined to verify proper placement of the injection and to determine the extent of local spread of the tracer. Only labelling from appropriately placed injections (15/16) was further examined. Next every third slide through the brain was inspected for the presence of red and green fluorescent Retrobeads on a wide field microscope (Nikon NiE equipped with an Andor Zyla 5.2. camera). Detailed notes of areas with retrogradely labelled cells were made on a copy of the Brain Atlas in Stereotaxic Co-ordinates (Paxinos and Watson, 1998).

Three animals in which the injection of Retrobeads was of very similar size and centred within the ICC at a midrostro-caudal level were chosen for quantitative analysis. For these three animals, low power confocal mosaic images of cortical regions and the IC injection sites were acquired with a Zeiss AxioObserver Z1, with a LSM800 confocal scan head, fitted with a motorised stage using a 20x air objective (0.8 numerical aperture (NA), 0.52 μs pixel dwell time, X-Y dimension 1024×1024, bit depth 8 with linear look-up tables (LUTs)). The images were examined and cells containing fluorescent beads were manually tagged in FIJI/ImageJ (Schindelin et al., 2012) using the Cell Counter plugin. Since the red beads were much more easily distinguished than the green beads, for quantification purposes, we counted only cells containing red beads in the cortices both ipsi- and contralateral to the injection site.

For **anterograde tracing experiments**, low-power confocal mosaic images (x20 air, 0.8 NA, 1024 pixels in the X–Y dimension, 0.52 μs pixel dwell time, bit depth 8 with linear LUTs) of the injection sites, were first acquired from DAPI labelled sections to verify the proper placement of injections. Low power confocal mosaic images (settings as above) of the whole IC (DAPI labelled sections) at a mid-rostro-caudal level were then acquired, and the presence of red and green dextran in the three subdivisions of the IC was examined. Since TRITC-dextran was more easily visualised and quantified than fluorescein-dextran, in six cases we examined only the distribution of red dextran in both hemispheres after injection into the right hemisphere (Fig. 5 a-d). In two cases (one visual cortex injection and one somatosensory cortex injection, Fig. 5 b and d) we examined the distribution of fluorescein-dextran after injection into the left hemisphere.

To determine whether dextran labelling in terminals was associated with GABAergic or non-GABAergic (putative glutamatergic) neurons, confocal Z-stacks of varying depths (7-15 μm) (x63 oil, 1.40 NA, 1.03 μs pixel dwell time, X-Y dimensions 1024×1024, Z step 0.26 μm, bit depth 8 with linear LUTs) were taken from the immunolabelled sections. Representative Z-stacks were taken in the ICD (2 stacks per animal), ICC (3 stacks per animal), and ICL (1 stack per animal). Z-stacks were cropped post-hoc to 7 μm thickness. The presence of red dextran was imaged in sections in which GABA had been labelled with AlexaFluor 488 (green) and green dextran was imaged in sections in which GABA had been labelled with AlexaFluor 568 (red).

Using Imaris™ (Bitplane) the DAPI signal was rendered to a ‘surface’ and the dextran labelling was rendered into ‘spots’. The proximity of the dextran ‘spots’ to DAPI ‘surface’ was measured, allowing us to assess whether dextran labelled terminals were in close proximity (≤ 3 μm) to cell somata. Note: we immunolabelled NeuN to visualise more of the cell body and proximal processes of the IC neurons to allow us to examine whether dextran labelled terminals contacted somatic or dendritic sites. However, we noted that in the IC, many DAPI labelled nuclei in the IC which were in close proximity to dextran labelled terminals and in some cases were also labelled for GABA, and so were presumed to be neurons, do not label for NeuN. Hence we used the DAPI nuclear signal rather than NeuN to define cell somata.

## RESULTS

### Retrograde labelling

#### Injection sites

We injected red Retrobeads into the right IC and green Retrobeads into the left IC of 8 rats (see Methods). Retrobead injection sites in all animals were inspected using a widefield microscope. At the injection site very bright beads could be seen. In all but one case Retrobead injections were well placed: one green Retrobead injection was misplaced and the data from this injection was discounted. Correctly placed injections (n=15) were centred in the ICC and included the lateral part of the ICD: in some cases there was some spread of Retrobeads into ICL. All injections were contained entirely within the boundaries of the IC. Notably, there was virtually no upward spread of Retrobeads into the visual cortex which overlies the IC. From the larger group of injections, the red Retrobead injections in three animals (R4, R5 and R6) were chosen for quantitative analysis. Figure 1 shows the extent of the red Retrobead injection in in the right IC in these animals traced from x20 confocal mosaic images.

**Figure 1:**
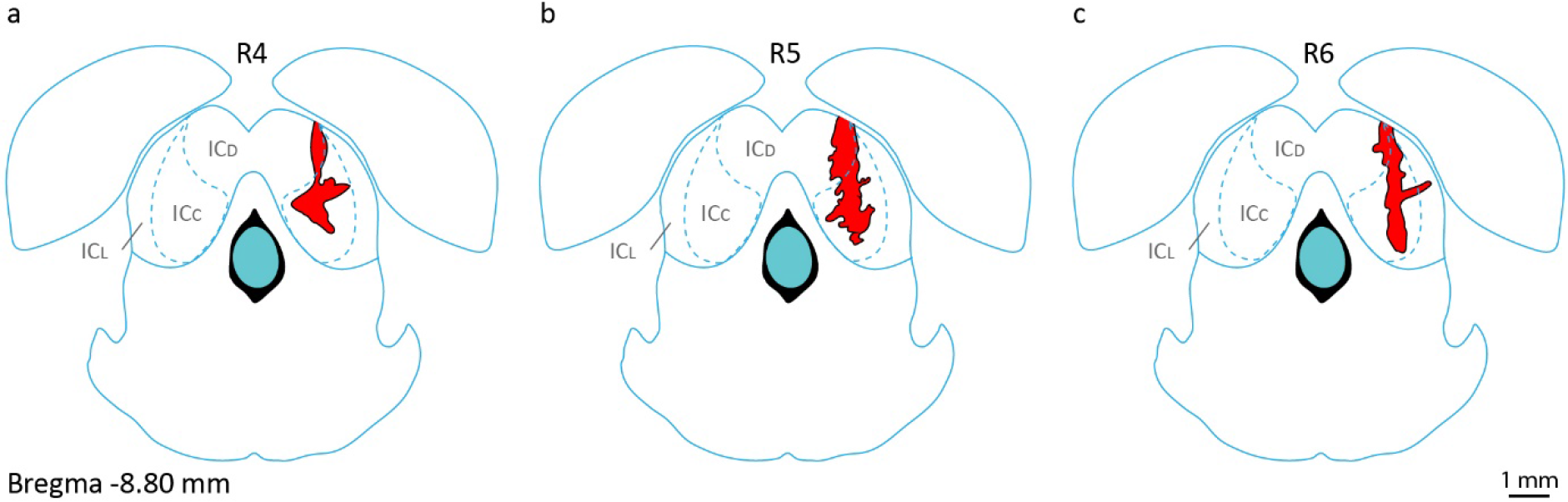
Sites of Retrobead injection in the right IC. For the three animals used for quantitative analysis (**a,** R4, **b,** R5, and **c,** R6), mosaic images were taken and the outline of each injection site was traced in Adobe Illustrator™ and superimposed on corresponding atlas outlines redrawn from Paxinos and Watson (1998). Figures show the rostro-caudal midpoint of the injection.

#### Retrograde labelling from the IC is seen in multiple cortical areas

As a positive control for our retrograde labelling, we first examined the presence of Retrobeads in the auditory cortex auditory cortex ipsilateral and contralateral to the injection site (henceforth referred to as the ipsilateral and contralateral cortices). As anticipated, Retrobeads injected in the ICC/ICD labelled many neurons in both the ipsilateral and contralateral auditory cortex, predominantly in layer V (Fig. 2a). Labelling was evident as distinct bright beads which are associated with DAPI stained nuclei suggesting that the beads are located within cell somata. The subcellular distribution of the Retrobeads and the orientation of the labelled cells indicates they are pyramidal neurons (Fig. 2b).

**Figure 2:**
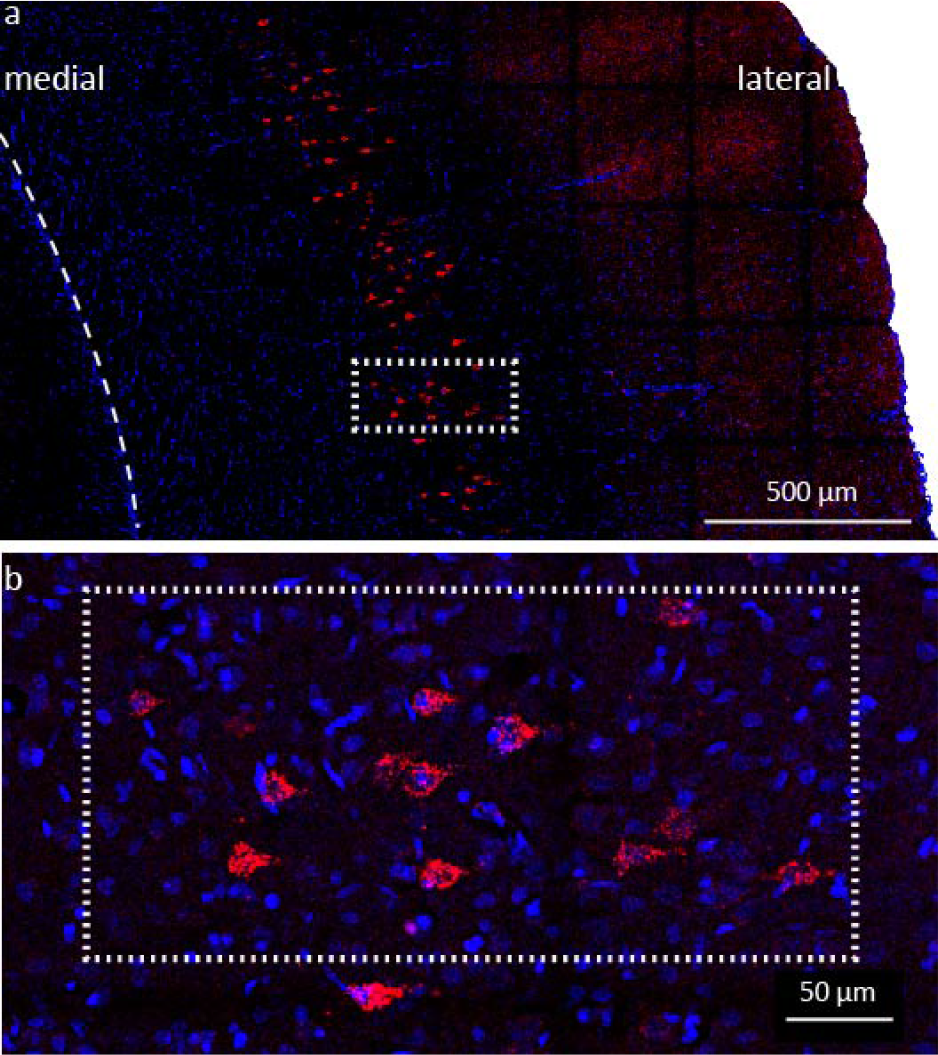
Neurons in the ipsilateral auditory cortex retrogradely labelled from the IC. Following injection of red Retrobeads into the right ICC/ICD, Retrobeads are found in neurons in the right auditory cortex. Nuclei are labelled with DAPI (blue). **a,** Retrogradely labelled cells are concentrated in a narrow band corresponding to layer V of auditory cortex. **b,** The distribution of beads in the somata highlights the pyramidal cell morphology of the retrogradely labelled neurons.

In addition to the labelling seen in the auditory cortex, large numbers of Retrobead-labelled neurons were also seen in many non-auditory cortical areas. Indeed, labelling extended from the visual areas caudally all the way to the prefrontal regions rostrally. Large numbers of neurons containing Retrobeads were found in visual cortex (Fig. 3a), somatosensory cortex (Fig. 3c), motor cortex (Fig. 3d and e) and prefrontal cortex (Fig. 3f). As in auditory cortex (Fig. 3b), Retrobeads were predominantly in the cell soma however, in comparison to the labelling in auditory cortex (Fig. 3b) in non-auditory cortical areas (Fig. 3a, c-f) the density of Retrobeads in each cell appeared lower.

**Figure 3:**
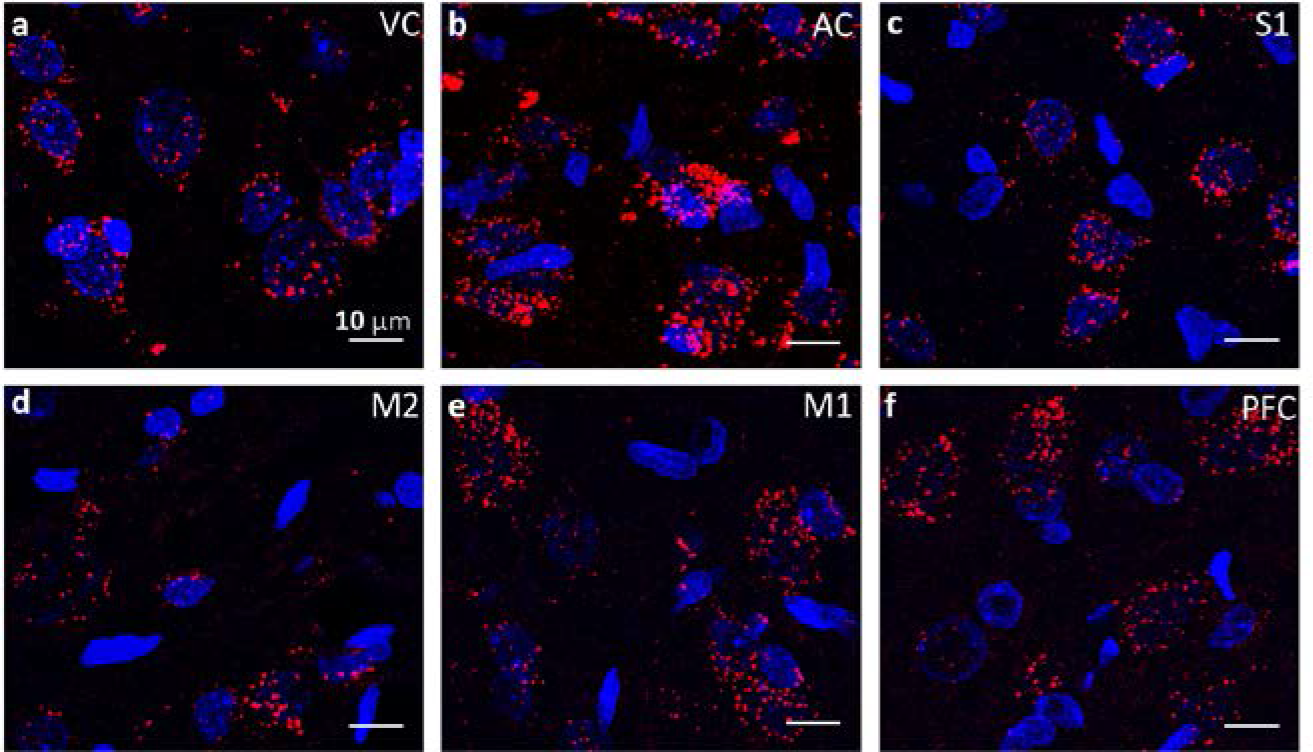
Multiple cortical regions are retrogradely labelled from the IC. High power maximum intensity projections from confocal Z-stacks showing red Retrobeads in neurons in cortical regions ipsilateral to the injection site. **a.** visual cortex (VC), **b.** auditory cortex (AC), **c.** primary somatosensory cortex (S1), **d.** secondary motor cortex (M2), **e.** primary motor cortex (M1) and **f.** prefrontal cortex (PFC). Nuclei are stained with DAPI (blue).

#### Retrograde labelling is present in both the ipsi- and contra-lateral hemispheres

##### Visual Cortex

Retrogradely labelled neurons were found in both ipsi- and contralateral visual cortices (Fig. 4a and b). They were numerous in both primary visual cortex (monocular and binocular) and secondary visual cortex (lateral area and mediolateral area) (Fig. 4a and b). Caudally, labelled cells were found in a wide band extending through layers III, IV and V (Fig. 4a) whereas more rostrally retrogradely labelled cells were concentrated in the deeper cortical layers (Fig. 4b). Quantification of the labelled neurons (in two sections from each of 3 animals) revealed a consistently high degree of lateralization. Thus, although retrograde labelling was bilateral, there were far fewer labelled neurons in the contralateral compared to the ipsilateral visual cortices (Fig. 4a, b and h).

**Figure 4:**
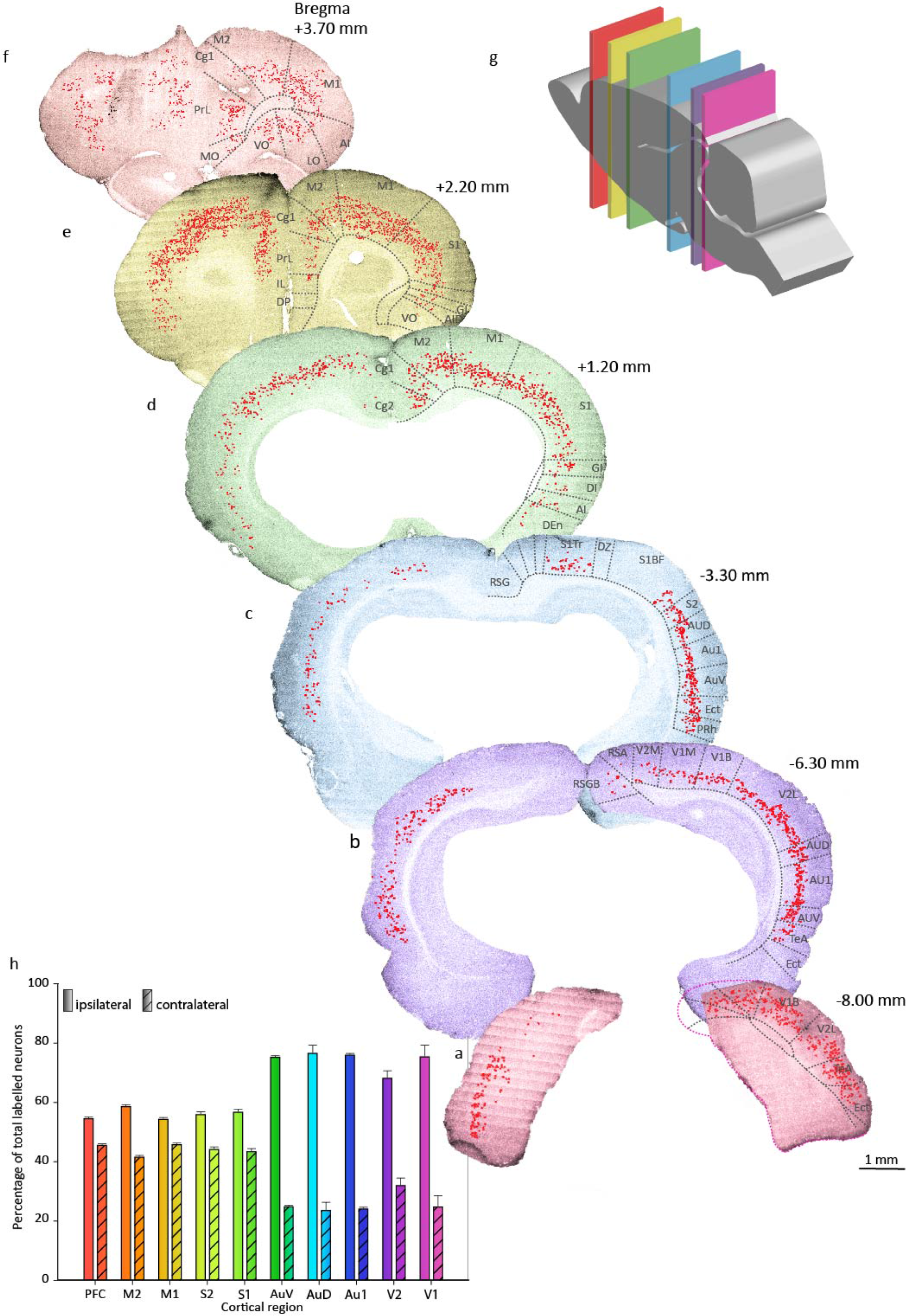
Retrograde cortical labelling demonstrates bilateral inputs to IC from multiple cortical regions. **a-f,** Confocal mosaic images of coronal cortical sections at multiple locations along the rostro-caudal axis (**g**) following the injection of red Retrobeads in the right ICC/ICD. Red markers indicate cells containing Retrobeads. Cortical divisions are redrawn from the atlas of Paxinos and Watson (1998) and region names are abbreviated on the sections. **h,** Proportions of retrogradely labelled cells in the cortex ipsi- and contralateral to the injected IC as percentage of total labelled neurons in each cortical region. Bars show mean + SEM (n=3 animals) derived from counts made in 2 or 3 sections per animal.

##### Auditory cortex

As expected (see Introduction), retrograde labelling was seen in both the ipsi- and contralateral auditory cortices (Fig. 4b and c). Many labelled neurons were present in the primary auditory cortex (Au1), and both dorsal (AuD) and ventral (AuV) secondary auditory cortices (Fig. 4b and c). On the ipsilateral side labelled neurons were densely packed in layer V with a few in layer VI whereas on the contralateral side, the labelled neurons were more scattered and occurred in layers III, IV and V (Fig. 4 b and c). As in the visual cortex, we observed many fewer retrogradely labelled neurons in the contralateral compared to the ipsilateral auditory cortex (Fig. 4h).

##### Somatosensory cortex

A second major non-auditory sensory cortical region which contained a substantial number of retrogradely labelled neurons was the somatosensory cortex. As was the case in the auditory and visual cortices, in the somatosensory cortex labelling was present in both ipsi- and contralateral hemispheres (Fig. 4c, d and e). Primary somatosensory regions (S1) with large numbers of retrogradely labelled cells included the trunk, forelimb, hindlimb, dorsal zone, and jaw areas. Note that although there are relatively few cells in the barrel field of case R5 (shown in Fig. 4c), R4 and R6 did have some labelled neurons in this region. Secondary somatosensory areas (S2) also contained large numbers of retrogradely-labelled cells (Fig. 4c). In more rostral sections the width of the band of labelled cells increases consistent with the increased thickness of the cortical layers (Fig. 4d and e). Interestingly, while there were more labelled neurons ipsilateral than contralateral to the injected IC, this difference was not as great as that observed in the auditory and visual cortices (Fig. 4h).

##### Motor cortex

Outside the sensory domain, we also found retrograde labelling from the IC in the motor cortex. Here, as in other cortical regions, labelling was evident in both ipsi- and contra-lateral sides. Many labelled neurons were seen in both the primary motor cortex (M1) and secondary motor cortex (M2) (Fig. 4d, e and f), and labelling extended along the whole rostro-caudal extent of the motor cortices. On both ipsi- and contralateral sides labelled neurons were dispersed through cortical layers III-VI. It was notable that in motor areas, the lateralization pattern of retrograde labelling resembled that seen in the somatosensory regions in that only marginally fewer labelled neurones were found in the contralateral compared to the ipsilateral side (Fig 4h).

##### Prefrontal cortex

The prefrontal cortex lies at the most rostral end of the cerebral cortex and comprises both the medial prefrontal and orbitofrontal regions. Following Retrobead injections into the IC many labelled neurons were found in the prefrontal cortex of both hemispheres (Fig. 3b, 4a-c).

The medial prefrontal cortex, which subserves executive function, is distinct in that its laminae lie in a dorso-ventrally. Retrograde labelled neurons were concentrated in the cingulate (Cg1 and Cg2) and prelimbic (PrL) cortices, and there were few, if any, labelled neurons in the more ventral infralimbic (IL) and dorsal peduncular (DP) regions. Retrogradely labelled neurons were located in the middle and deep layers (III-VI) close to the white matter of the forceps minor.

The orbitofrontal cortex, which is situated ventral to the forceps minor, integrates sensory and autonomic information. In orbitofrontal cortex, retrograde labelled neurons were observed in the ventral and lateral regions with fewer labelled neurons in the medial orbitofrontal cortex. Retrograde labelled neurons were located in the deeper layers.

Considering the prefrontal cortex as a whole, the degree of lateralization was low with almost as many retrogradely labelled neurons on the contralateral side (44%) as on the ipsilateral side (56%) (Fig 4h).

##### Non-cortical regions are retrogradely labelled from IC

Following Retrobead injections into the IC, labelled cells were also observed in brainstem auditory regions known to innervate the IC (including cochlear nucleus, olivary complex, dorsal nucleus of the lateral lemniscus). In addition, we noted Retrobead-labelled cells in many subcortical forebrain regions including olfactory bulb, caudate putamen, hippocampus, thalamus, hypothalamus and amygdala, as well as in mid- and hindbrain regions including superior colliculus and dorsal raphe (Olthof et al., 2019). The innervation of the IC by these regions will be the subject of a second paper.

Although we concentrated our detailed analysis on the red Retrobeads and quantified retrograde labelling in only a subset of the animals, qualitatively the distribution of retrogradely labelled cells was the same for all red and green Retrobead injections.

##### Anterograde labelling

To verify our findings with retrograde tracing, and to examine both the regional distribution and neuronal targets of cortical inputs to the IC, we injected anterograde tracers (TRITC- (red) and fluorescein- (green) labelled dextran) into cortical regions of interest and examined the presence of dextran labelled terminals in the ipsi- and contralateral IC.

#### Cortical sites of anterograde tracer injections

At the injection site, fluorescent labelling of neuronal elements was intense and the location and dispersal of the tracer was readily visualised. All injections were correctly placed within the intended cortical region of interest. Although most animals had two cortical injection sites (one red (right) one green (left)), we selected just one of these sites in each animal. As a result, labelling originating from one prefrontal and one motor cortex injection, and two auditory, two visual, and two somatosensory cortical injections were studied in detail (Fig. 5a-d).

**Figure 5:**
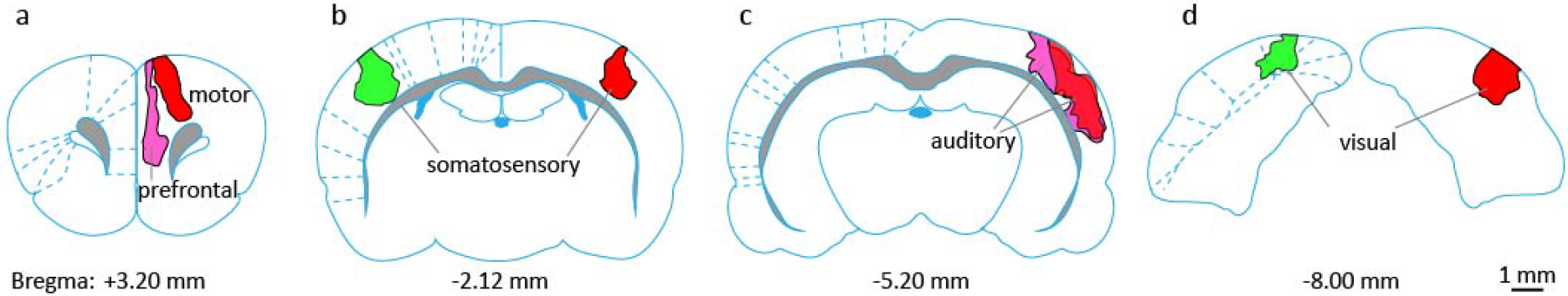
Sites of anterograde tracer injection in the cortices. TRITC (red) labelled dextran anterograde tracer was injected in the right hemisphere and fluorescein (green) labelled dextran in the left hemisphere. For the injection sites selected for analysis, mosaic images were taken and the outline of each injection site was traced in Adobe Illustrator™ and superimposed on corresponding atlas outlines redrawn from Paxinos and Watson (1998). Figures show the rostro-caudal midpoint of the injection. Note: each area traced is from a different animal. Injection sites in **a,** prefrontal cortex (Cg1 and PrL, magenta) and motor cortex (M2, red), **b,** somatosensory cortex (S1, red and green), **c,** auditory cortex (Au1, AuD, AuV, PtA, red and magenta), and **d,** visual cortex (V1, green and V1 and V2, red).

#### Anterograde labelling confirms non-auditory cortical projections to IC

For each of the cortical regions of interest highlighted by our retrograde findings (visual, auditory, somatosensory, motor, and prefrontal), injection of dextran resulted in dense anterograde labelling (see Fig. 5c). Dextran labelling was evident as individual small fluorescent puncta or clusters of puncta (presumed neuronal terminals) and was present in all divisions of the IC (ICD, ICC, and ICL). We calculated the distance of the dextran labelled terminals from DAPI labelled nuclei using the Imaris™ (Bitplane) ‘spot’ and ‘surface’ functions. This analysis revealed that the majority of dextran labelled terminals were clustered around DAPI labelled cell nuclei, sometimes apparently extending along proximal processes but within 3 μm of DAPI labelled nuclei (Fig. 6 top row). Other labelled terminals were further away from cell somata, presumably on other neuronal elements (Fig. 6). This pattern of labelling was seen irrespective of the cortical site of injection of the tracer or the subdivision of the IC examined.

**Figure 6:**
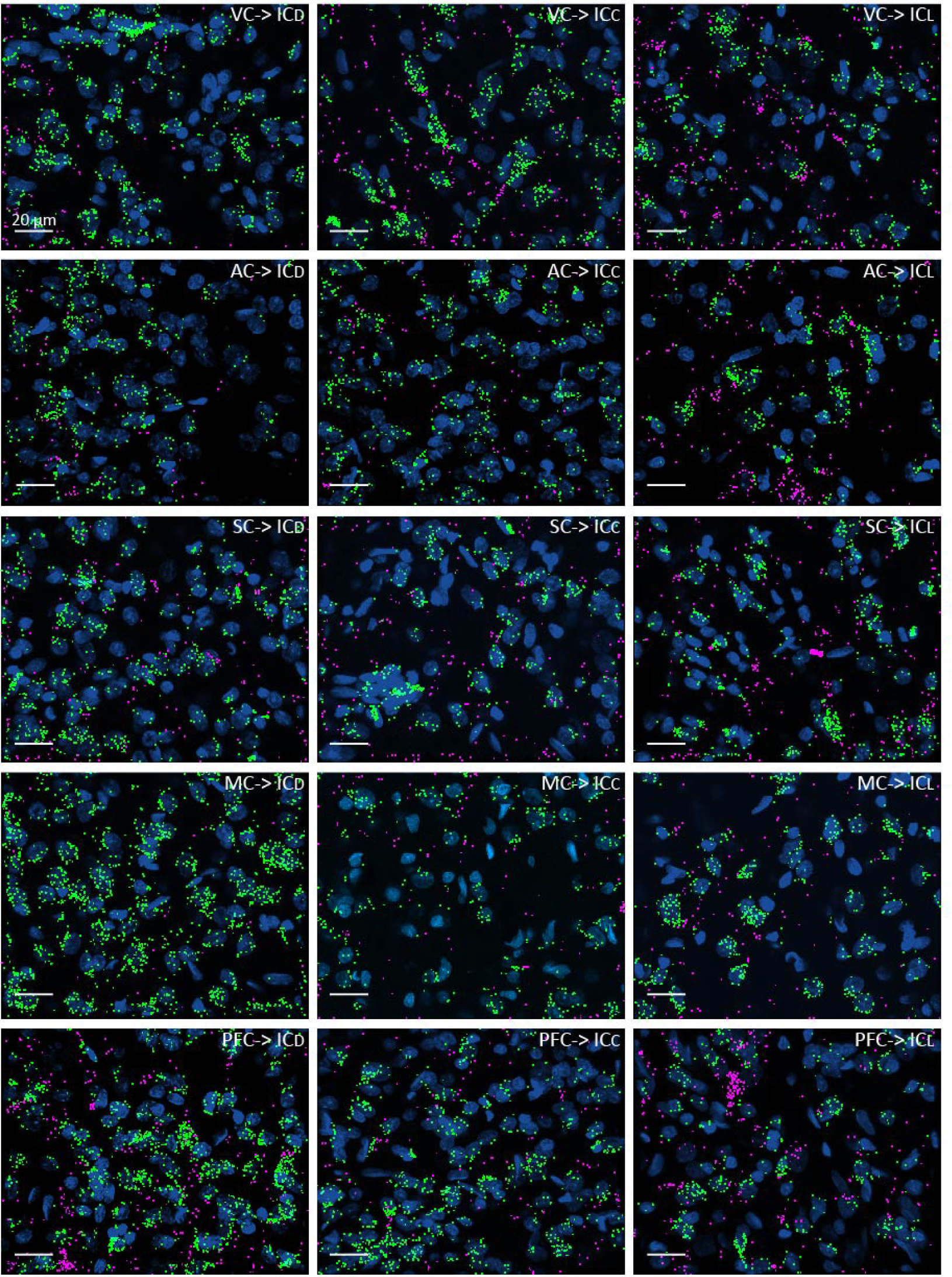
Anterograde tracing reveals terminals originating from multiple cortical regions target neurons in all IC subdivisions. Anterograde dextran labelling and were converted into ‘spots’ and DAPI stained nuclei were converted to ‘surfaces’ (blue) using Imaris™ and the proximity of spots to surfaces was determined. Spots closer than 3μm are colored green and those further away are colored magenta. Distribution of anterograde labelled terminals in ICD, ICC, and ICL (columns left to right) following injections of tracer into visual, auditory, somatosensory, motor and prefrontal cortices (rows top to bottom).

Our retrograde labelling showed that the largest difference in the lateralisation of cortico-collicular neurons was between the auditory cortex (24% contralateral) and motor cortex (46% contralateral). Hence, for these two regions, we compared anterograde labelling in the IC ipsi and contralateral to the injection (Fig. 7). For both auditory cortex (Fig. 7a) and motor cortex (Fig. 7b) tracer injections, dextran labelled terminals were observed in both ICs. Counts of dextran spots in five Z-stacks (one in ICD and ICL, and 3 in ICC) obtained from one animal on both the ipsi- and contralateral side confirmed the findings from the retrograde studies, there was evidence for highly lateralised connections from auditory cortex (62.4 ± 2.12 % ipsi- vs. 37.6 ± 1.48 contralateral) contrasting with strongly bilateral connections from the motor cortex (51.5 ± 1.47% ipsi- vs. 49.5 ± 1.97 contralateral).

**Figure 7:**
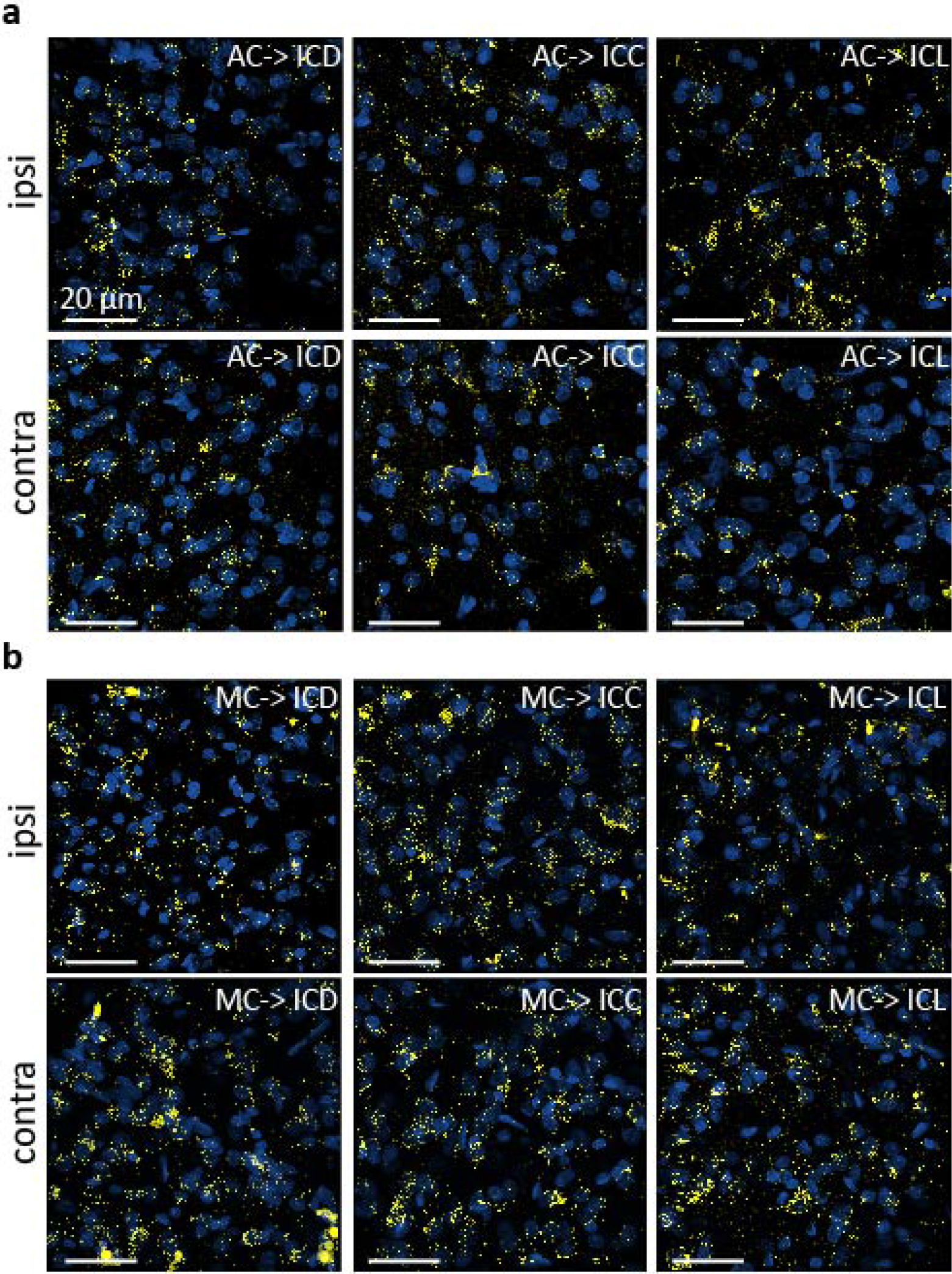
Anterograde labelling is present in the ICs both ipsi- and contralateral to the cortical injection. Anterograde dextran labelling (yellow) in the three subdivisions of the IC (ICD, ICC, and ICL). Dextran injected in **a,** auditory cortex (AC, see Fig 4c). **b,** motor cortex (MC, see Fig4a). Cell nuclei are stained with DAPI (blue).

#### Both GABAergic and glutamatergic neurones are targeted by cortical inputs

In sections from our anterograde tracing studies, immunolabelling for GABA revealed a high density of GABAergic neurons of various sizes and morphologies in the IC. Regardless of the cortical site at which the dextran tracer was injected, labelled terminals were found in close proximity to many large (> 15 μm) (Fig. 8ai-iii) as well as smaller (Fig. 8bi-iii) GABAergic neurons. In the ICL in particular, we saw GABA neurons with extensive dendrites, and dextran labelled terminals were frequently found in close proximity to these structures (Fig. 8ciii). Labelled terminals were also found close to the smaller, finer, GABAergic dendrites seen in ICD and ICC (Fig. 8ci and cii). We noted labelled terminals close to some DAPI stained nuclei that did not label for GABA. Since it is well established that IC neurons contain only one of two principal neurotransmitters, glutamate or GABA (Merchán et al., 2005), these are presumed to be glutamatergic neurons (Fig. 8di-iii).

**Figure 8:**
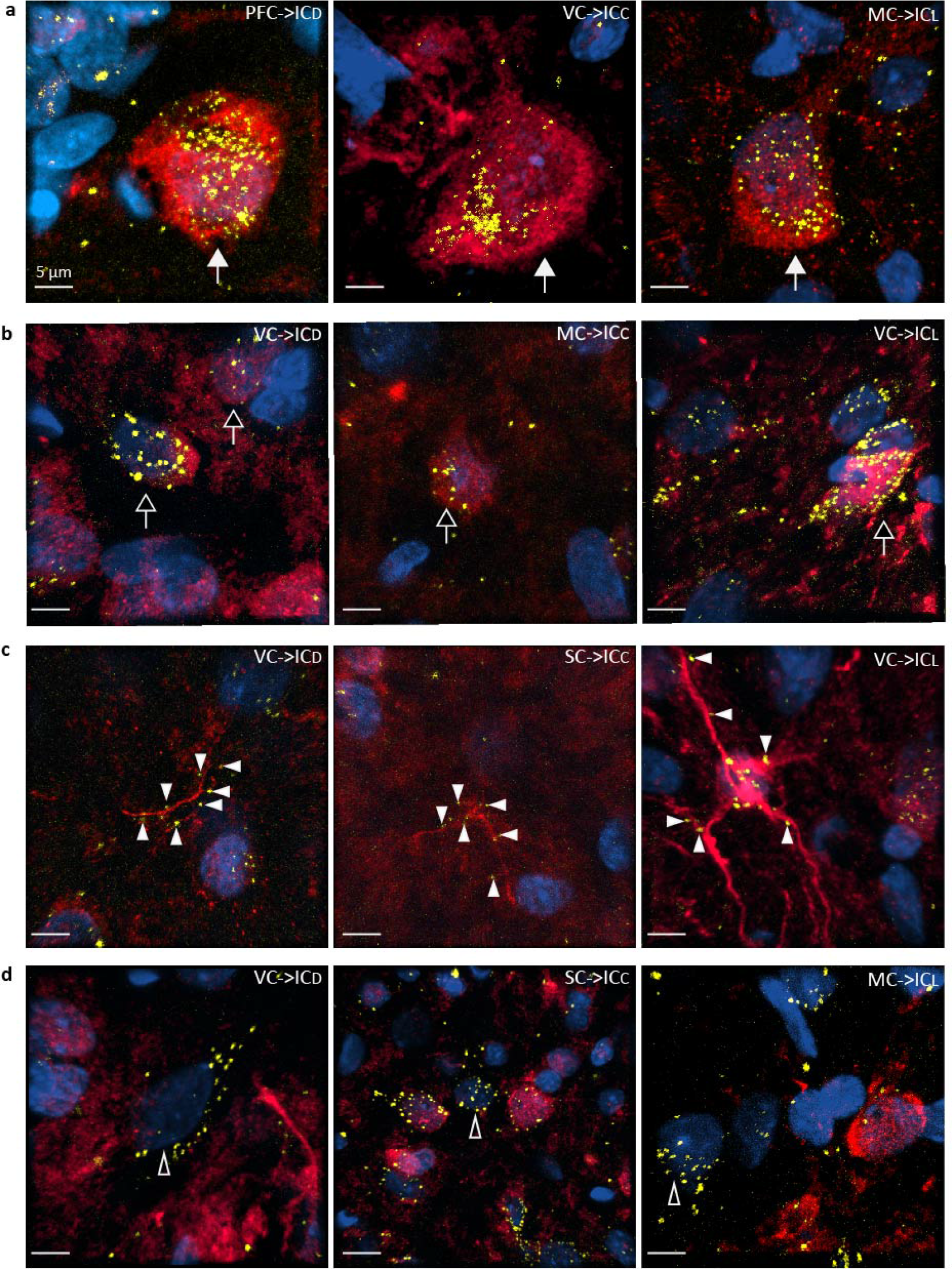
Terminals originating from multiple cortical regions associate with both GABAergic and glutamatergic neurons in the IC. Confocal Z-stack images showing dextran labelled terminals (yellow) and DAPI stained nuclei (blue). Terminals originating from cortical regions are evident in close proximity to **a,** large GABAergic neurons (red, closed arrows), **b,** smaller GABAergic cells (red, open arrows), and **c,** GABAergic dendrites (red, closed arrowheads). **d,** terminals of cortical origin are also found in close proximity to neurons which do not label for GABA-presumed glutamatergic neurons (open arrowheads). Example confocal images from ICD, ICC, and ICL (columns left to right) showing dextran labelling following injection into visual, somatosensory, motor, or prefrontal cortices (see individual panel labels).

## DISCUSSION

The most important finding arising from our retrograde and anterograde tracing studies is that multiple regions of the cerebral cortex make extensive bilateral connections with all three subdivisions of the IC. In addition to previously established cortico-collicular inputs from the auditory cortex we demonstrate substantial connections from the visual, somatosensory, motor and prefrontal regions. Retrograde tracing shows that connections from the ipsilateral side are the most abundant. However, the ipsi-contralateral difference is much less pronounced for the somatosensory, motor and prefrontal cortices where numbers of labelled neurons are nearly equal, than for the visual and auditory cortices where the difference is approximately threefold. In the main, cortico-collicular neurons are most abundant in layer V of the cortex. Anterograde tracing suggests that terminals of fibers from these cortical areas form synapses with both GABAergic and putative glutamatergic neurons in all three divisions of the IC. A surprising aspect of our results is the extent to which diverse cortical projections target the ICC, this subdivision is the main recipient of ascending input from brainstem auditory nuclei and traditionally regarded primarily as a centre of auditory processing (Oliver, 2005; Ito and Malmierca, 2018).

### Technical considerations

The combined application of retrograde and anterograde tracing methods provides convincing evidence for the existence of cortico-collicular connections from non-auditory cortical regions. As demonstrated in previous studies, Retrobeads show minimal diffusion (Katz et al., 1984) and our injections which targeted ICC and ICD resulted in minimal spread into ICL. Importantly, no Retrobeads were observed in the overlying visual cortex. Thus, we can be confident that the cortical labelling we observed resulted from retrograde transport from the IC. Although we quantified data using red Retrobeads from three exemplar animals, we saw the same pattern of labelling with both red and green Retrobeads in all eight animals.

There were differences in the density of Retrobead labelling within neurons, both within and between regions. Some somata contained very large numbers of beads while others, often close by, contained far fewer beads. It was notable that neurons in the ipsilateral auditory cortex contained the highest number of very densely labelled cells. Retrobeads are considered efficient tracers, nevertheless differences in the density of labelling of neighbouring cortical neurons have been reported previously (Schofield et al., 2007). Such differences in the density of retrograde labelling may relate to the total number, or the pattern, of terminal distribution in the IC, or indeed to some other factor such as the level of neuronal activity. Differences in labelling density are unlikely to be explained by the distance from the IC to the soma as both densely and less densely labelled cells were found juxtaposed to one another in all cortical regions.

We chose a relatively high molecular weight dextran (10,000 m.wt) conjugated to a fluorophore, in order only to label terminals. Labelling terminals in this way avoids the high background labelling that results from the labelling of other neuronal elements. Anterograde tracing shows that cortico-collicular projections from non-auditory cortices target all three subdivisions of the IC. Because we did not make multiple, systematically placed injections in each of the cortices studied, we cannot comment on the relative abundance of cortico-collicular connections to the different subdivisions of the IC; such an analysis will require further study.

### Auditory and non-auditory cortico-collicular projections

We found evidence for a dense and highly lateralised projection from the auditory cortex to the subdivisions of the IC. Feedback connections from the auditory cortex to the inferior colliculus were first reported by (Beyerl, 1978) in rat and have since been described originating from multiple fields of the auditory cortex in several species (see (Winer, 2005; Schofield, 2010; Bajo and King, 2013) for reviews). While cortico-collicular projections to the cortical subdivisions of the IC are known to be particularly dense, connections to central nucleus have also been reported in rat (Feliciano and Potashner, 1995; Saldana et al., 1996) and other species (Budinger et al., 2000; Bajo et al., 2007). Furthermore, a tonotopic correspondence between the frequency representations in the cortex and the ICC has been demonstrated both anatomically (Andersen et al., 1980; Saldana et al., 1996; Bajo and Moore, 2005; Bajo et al., 2007) and functionally (Lim and Anderson, 2007). Our findings, confirming these previous studies, thus serve as a positive control for our exploration of non-auditory cortico-collicular connections.

We demonstrate extensive bilateral cortico-collicular projections from visual, somatosensory, motor and prefrontal cortices. Although ours is the first study to reveal the full extent of these connections, evidence pointing to the existence of non-auditory cortico-collicular connections can be found in an earlier degeneration study in cat by Cooper and Young (1976). Following large lesions of the visual, somatosensory or motor cortices, these authors reported degenerating fibres in the ipsilateral IC. Their paper provides no quantification to allow the extent of this degeneration to be assessed, but their figures suggested these connections are sparse and mostly confined to areas equivalent to ICD and ICL. More recently, Lesicko et al. (2016), using anterograde tracing, demonstrated connections from the somatosensory cortex to the IC in mouse, but they only reported on the presence of labelled terminals in the so-called ‘neurochemical modules’ in ICL (Chernock et al., 2004). These modules also receive ascending somatosensory inputs from the dorsal column nuclei and spinal trigeminal nucleus consistent with physiological evidence for ascending somatosensory input to ICL (Aitkin et al., 1978; Aitkin et al., 1981; Jain and Shore, 2006). In our study, we observed connections distributed more widely, both within ICL and throughout the other IC subdivisions.

In the visual modality we observed extensive cortico-collicular connections from both V1 and V2 to the IC. Several functional studies in IC have reported visual modulation of neuronal responses to sounds, and even neuronal responses to visual stimuli alone (Syka and Radil-weiss, 1973; Tawil et al., 1983; Mascetti and Strozzi, 1988; Porter et al., 2007; Bulkin and Groh, 2012). In the cortical subdivisions these interactions may be explained by direct retinal input to these regions (Itaya and Van Hoesen, 1982; Yamauchi and Yamadori, 1982; Herbin et al., 1994). However, the longer latency of responses observed by Porter et al. (2007) are consistent with effects mediated by indirect pathways via the superior colliculus (Adams, 1980; Coleman and Clerici, 1987), or from the visual cortical input we describe here.

The existence of connections from the visual cortex to the IC may explain recent functional studies in which the BOLD response recorded in the IC using fMRI was influenced by ablating the visual cortex, or by modulating visual cortical output using optogenetics (Gao et al., 2015; Leong et al., 2018). Interestingly, the results of these studies suggest that inputs from visual cortex enhanced response gain, whereas auditory cortical inputs reduced response gain and increased response selectivity to species-specific vocalisations. Although cortical output projections are believed to be exclusively excitatory, our finding that projections from cortical regions terminate on both GABAergic and presumed glutamatergic neurons shows that cortically mediated excitation can readily be transformed to inhibition in the IC by such di-synaptic circuits.

The feedback connections from motor cortex to the inferior colliculus we report here may be important for two reasons. First, they are a source of feedback about self-generated sounds. Such sounds could include self-generated vocalisations as well as the sounds generated when an animal interacts with its environment (Schneider and Mooney, 2018; Schneider et al., 2018). The ability to distinguish self-generated sounds from those originating independently of it, is critical for an animal’s survival. Motor and somatosensory cortical feedback could provide predictive information that enables sounds generated by vocalisations or other movements to be discounted during the process of parsing of simultaneous sound sources. The fact that neurons throughout the motor cortex project to the IC suggests that all body movements may contribute to this process, and that the feedback is not limited to the control of vocalisation. A second reason why motor inputs to the IC are significant is that the IC constitutes part of a network of brain centres involved in escape and other fear responses to aversive sounds. Blockade of inhibition and modulation of NMDA receptors in the IC can trigger such behaviours (Brandão et al., 1988; Cardoso et al., 1994; Nobre et al., 2004) and auditory responses in IC are enhanced by unconditioned and conditioned fear evoking stimuli (Brandao et al., 2001).

We know of no existing evidence that the prefrontal cortex influences responses in the IC. However, neurons in the orbitofrontal regions respond to sound and via descending connections influence sound processing in auditory cortex (Saunders et al., 1985; Fritz et al., 2010; Schneider et al., 2014; Schneider et al., 2018; Winkowski et al., 2018). These prefrontal cortical regions of cortex have a central role in executive function and exert task-dependent influences on sensory processing (Öngür and Price, 2000). Our results demonstrate direct projections from the medial and orbital prefrontal cortex to the IC and suggest that higher-level cognitive functions, including goal directed behaviour, directly affect auditory processing in the midbrain.

Interestingly, whereas the numbers of neurons projecting to the IC from the visual and auditory cortices displayed a marked ipsilateral dominance, those in the somatosensory, motor, and prefrontal cortices were more equal in number between the two sides. This might reflect the fact that movement elicited sounds and cognitive mechanisms are not as strongly lateralised as sources of auditory and visual signals.

The extensive descending input to the IC from several distinct areas of the cerebral cortex leads us to conclude that the IC is much more than an integrator of auditory signals ascending from the brainstem. The diversity of descending inputs to this midbrain structure suggest it has a complex, nuanced role in auditory perception. It demonstrates that multisensory influences from the cortex operate at an earlier stage than previously believed. We suggest that this multisystem cortico-collicular information plays a fundamental role, perhaps through predictive coding, in an animal’s perceptual and behavioural responses to sound. These findings also emphasise that a complete understanding of the role of the IC requires studies in awake, behaving animals in which these descending, sensory, motor, and cognitive signals can make their full contribution to IC function.

The extensive descending input to the IC from several distinct areas of the cerebral cortex leads us to conclude that the IC is much more than an integrator of auditory signals ascending from the brainstem. The diversity of descending inputs to this midbrain structure suggest it has a complex, nuanced role in auditory perception. These findings also demonstrate that multisensory influences from the cortex operate at an earlier stage of processing than previously believed. We suggest that this multisystem cortico-collicular information plays a fundamental role, perhaps through predictive coding, in an animal’s perceptual and behavioural responses to sound. These findings also emphasise that a complete understanding of the role of the IC requires studies in awake, behaving animals in which these descending, sensory, motor, and cognitive signals can make their full contribution to IC function.

As discussed in the Introduction, the traditional view of sensory processing as both centripetal and unimodal has recently been challenged. It is now established that there are both corticofugal connections allowing top down modulation and cortico-cortical connections allowing interactions between sensory modalities. The present data indicate that there are further pathways to subcortical sensory centers from cortical regions subserving not only sensory, but also motor and executive function.

## LIST OF ABBREVIATIONS

AC: auditory cortex
AI: agranular insular cortex
Au1: primary auditory cortex
AuD: secondary auditory cortex dorsal zone
AuV: secondary auditory cortex, ventral zone
Cg1: cingulate cortex area 1
Cg2: cingulate cortex area 2
Den: dorsal endopiriform nucleus
DI: dysgranular insular cortex
DP: dorsal peduncular cortex
DZ: primary somatosensory cortex dorsal zone
Ect: ectorhinal cortex
GI: granular insular cortex
IC: inferior colliculus
ICC: central nucleus of inferior colliculus
ICD: dorsal cortex of inferior colliculus
ICL: lateral cortex of inferior colliculus
IL: infralimbic cortex
LO: lateral orbital cortex
M1: primary motor cortex
M2: secondary motor cortex
MO: medial orbital cortex
PFC: prefrontal cortex
Prh: perirhinal cortex
PrL: prelimbic cortex
RSA: retrosplenial agranular cortex
RSG: retrosplenial granular cortex
RSGb: retrosplenial granular b cortex
S1: primary somatosensory cortex
S1BF: primary somatosensory barrel cortex
S1Tr: primary somatosensory cortex trunk region
S2: secondary somatosensory cortex
TeA: temporal association cortex
V1B: primary visual cortex, binocular area
V1M: primary visual cortex, monocular area
V2L: secondary visual cortex, lateral area
V2M: secondary visual cortex, mediolateral area
VC: visual cortex
VO: ventral orbital cortex

## ACKNOWLEDGEMENTS

This work was supported by the BBSRC (Grant BB/P003249/1 to A.R. and S.E.G.). We thank the Newcastle University BioImaging Unit for their support and assistance in performing the microscopy. Andrew Trevelyan for advice on injecting tracers.

## DATA AVAILABILITY STATEMENT

The datasets generated during and/or analysed during the current study are available from the corresponding author on reasonable request

## AUTHOR CONTRIBUTIONS

BMJO, AR and SEG all contributed to the design of the experiments and the injection of tracers. BMJO and SEG processed and BMJO imaged the resulting tissue. BMJO, AR and SEG wrote and edited the manuscript.

